# Human foot force reveals different balance control strategies between healthy younger and older adults

**DOI:** 10.1101/2024.04.08.588635

**Authors:** Kaymie Shiozawa, Rika Sugimoto-Dimitrova, Kreg G. Gruben, Neville Hogan

**Affiliations:** Department of Mechanical Engineering, Massachusetts Institute of Technology, Cambridge, MA 02139, USA; Department of Mechanical Engineering, University of Wisconsin – Madison, Madison, WI 53706, USA; Department of Kinesiology, University of Wisconsin – Madison, Madison, WI 53706, USA; Department of Brain and Cognitive Sciences, Massachusetts Institute of Technology, Cambridge, MA 02139, USA

**Keywords:** aging, balance, ground reaction force, inverted pendulum, neural control

## Abstract

Aging can cause the decline of balance ability, which can lead to increased falls and decreased mobility. This work aimed to discern differences in balance control strategies between healthy older and younger adults. Foot force data of 38 older and 65 younger participants (older and younger than 60 years, respectively) were analyzed. To first determine whether the two groups exhibited any differences, this study incorporated the orientation of the foot-ground interaction force in addition to its point of application. Specifically, the frequency-dependence of the “intersection point” of the lines of actions of the foot-ground interaction force were evaluated. Results demonstrated that, like the mean center-of-pressure speed, a traditionally-employed measure, the intersection-point analysis could distinguish between the two participant groups. Then, to further explore age-specific control strategies, simulations of standing balance were conducted. An optimal controller stabilized a double-inverted-pendulum model with torque-actuated ankle and hip joints corrupted with white noise. The experimental data were compared to the simulation results to identify the controller parameters that best described the human data. Older participants showed significantly more use of the ankle than hip compared to younger participants. Best-fit controller gains suggested diminished intrinsic muscle stiffness in older adults, indicative of muscle strength loss, that was likely compensated by increased neural feedback. These results underscore the advantages of the intersection-point analysis to quantify shifts in interjoint control strategies with age, thus highlighting its potential to be used as a balance assessment tool in research and clinical settings.

**NEW & NOTEWORTHY:** Age groups were distinguished by analyzing foot-ground force data during quiet standing in older and younger adults to calculate the foot-force vector intersection point that emerges across frequency bands. Modeling balance and comparing the simulations’ outcomes with experimental results suggested that older adults increased reliance on neural feedback, possibly compensating for muscle strength deficiency. This novel analysis also quantified controllers for each participant, highlighting its potential to be implemented as a balance assessment tool.

## INTRODUCTION

Advancing age is associated with neuro-physiological changes that challenge the human neuromotor control system. Weakened skeletal muscles, reduced proprioception, and slowed nerve transmission speed are some deficits that can severely compromise the ability to control movement (1–5). One major consequence is deterioration of balance control, marked by increase in falls (6, 7). Unintentional falls are the predominant cause of injuries among older adults and contribute to an average of 100 deaths per day in the United States alone (8). Furthermore, postural control deficits tend to reduce bipedal mobility, degrading capacity for independent living (9). Thus, there is a pressing need to understand the effects of aging on balance impairment to mitigate fall risk and preserve mobility among older adults.

The neuromuscular mechanisms underlying human posture control remain poorly understood. Prior research on assessing balance often involved the use of external perturbations to probe the underlying neuromuscular balance controller (10). Though useful as a controlled experimental paradigm, such “reactive balance” tests often require specialized equipment to implement the perturbations and could involve artificial settings that may not transfer well to balance control in everyday life (10). Additionally, for some older adults who are already at risk of falling, perturbations increase injury risk.

On the other hand, perturbation-free balance assessments make use of voluntary movements and intrinsic variability in the neuromotor control system to investigate the balance controller in a more natural and safe setting. The majority of prior quiet-standing studies on older adults used a singleinverted-pendulum model, which focused on the contribution of the ankle joint (10). However, recent work showed that the inverted-pendulum model is insufficient to capture variations in the neuromotor control strategy inferred from foot-ground interaction forces (11, 12). An augmented model of quiet-standing balance, with more degrees of freedom, should provide further insight into the age-related differences in posture control.

Prior work on balance has also largely focused on measurements of postural sway, either by tracking the center-of-pressure trajectory under the feet via a force plate or the center-of-mass trajectory estimated from motion capture. Maki et al. and Prieto et al. studied various center-of-pressure-based measures extensively and identified mean center-of-pressure speed as the measure that best distinguished older and younger adults (13, 14). Subsequent work has attempted to develop mathematical models to predict differences in the control mechanisms that explained the variations in postural behavior observed with age (15, 16).

However, confining analyses to only center-of-pressure trajectory and variability neglects other components of the foot-ground interaction force vector that are both critical to the physics of the task and reflect neuro-motor control strategies (14, 17). Recent work suggest that incorporating the orientation of the foot-ground interaction force in addition to its point of application (center-of-pressure position) offers insight into the dynamics and control of balance (11, 18). Specifically, a measure termed the “intersection-point height” was developed to quantify emergent relative behavior of these two signals (18). Experiments revealed that the center-of-pressure position and foot-force orientation covary with a ratio dependent on frequency (18). Subsequent numerical analysis based on a double-inverted-pendulum model indicated that the measure reflects distinct control strategies (11). Consistent with that result, empirical studies showed that support surface properties, muscle contraction state, and stroke also systematically affect the measure (12, 19–21). The variation of the frequency-dependent intersection point under different balance conditions suggests its potential utility for understanding agerelated differences in balance ability.

In this work, we investigated how the balance strategies of older adults differ from those of younger adults. We hypothesized that: 1) the intersection-point-height measure can distinguish between older and younger participants, and 2) a mathematical model that reproduces the intersection-point-height data can inform age-related differences in balance strategies.

To test these hypotheses, we used existing data of 38 unimpaired older participants and 65 unimpaired younger participants, who stood quietly with feet side-by-side on a force plate (22). The sagittal-plane data of the foot-ground interaction force were processed to examine the frequency-dependent intersection-point height. The mean anteroposterior center-of-pressure speed was computed from the same dataset for comparison with the intersection-point measure. Then, assuming that a quietly standing human can be adequately modeled using a double-inverted pendulum with torque-actuated ankle and hip joints corrupted by white noise, an optimal control method was employed to simulate balance. Simulated data were compared to human data, and control parameters that resulted in the best-fit of the average and individual participant data for both age groups were reported. Stiffness gain matrices were also computed based on these best-fit parameters.

Results showed that both the intersection-point measure and the conventionally-used center-of-pressure speed were able to distinguish between age groups. Model-based analysis of the intersection point results quantified age-specific control strategies: older adults penalized the hip joint significantly more than younger adults. Optimal feedback gain values suggested increased dependence on neural feedback in older adults to potentially compensate for decreased intrinsic muscle stiffness. These results underscore the advantages of the intersection-point analysis to quantify age-related variations in control strategies.

## MATERIALS AND METHODS

### Human Experiment

The participants’ age and posturography information from the public data set by Santos and Duarte (22) were used for this study. Details of the experiment beyond what is outlined below can be found in a previous study (22).

### Participants

Out of the 163 participants who were recorded in the public data set (22), data from 60 participants who reported a condition that could have affected their ability to stand were omitted. Thus, data from 103 participants were used in this study; 38 participants (10 males and 28 females) were 60 years and older, while 65 participants (27 males and 38 females) were under 60 years old. The older participants’ average age, mass, and height were 70.7 ± 6.7 years, 63.1 ± 8.1 kg, 1.57 ± 0.09 m, respectively, while the younger participants’ average age, mass, and height were 26.9 ± 7.3 years, 62.6 ± 7.8 kg, 1.68 ± 0.08 m, respectively (mean ± standard deviation).

### Protocol & Apparatus

Participants were asked to stand barefoot on a rigid horizontal force plate surface with their heels 10 cm apart and their feet angled at 20 degrees off the mid-sagittal plane. They were tasked to stand as still as possible with their arms at their sides while gazing at a point marked on the wall (3 m away). Three trials, each lasting 60 s, were conducted under this condition for each participant. Foot-force data were collected at 100 Hz using a force plate (OPT400600-1000; AMTI, Watertown, MA, USA); the data were subsequently low-pass-filtered down to 10 Hz with a 4th order zero-lag Butterworth filter (22).

## Data Analyses

The intersection-point height across various frequencies were computed by processing the force data to determine whether the analysis could distinguish between older and younger participants. This method was compared against a traditionally employed measure, mean center-of-pressure speed.

### Intersection-Point Analysis

Previous work defined the intersection point as the point that minimally deviated from bandpass-filtered sagittal-plane foot-force vectors (18). To compute the intersection point, it was assumed that variations in the foot-force orientation were small to allow the foot-force orientation (θ_*F*_) to be approximated as

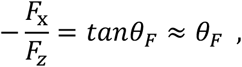

where *F*_x_ and *F*_*z*_ are the horizontal and vertical components of the foot-force vector (***F***). Fig. 1 depicts these signals. Both θ_*F*_ and the center-of-pressure position (*x*_*CoP*_) were analyzed in the frequency domain to investigate the system response. To conduct this frequency-domain analysis, a spectral-based method was employed as outlined by Sugimoto-Dimitrova et al. (23).

**Figure 1.**
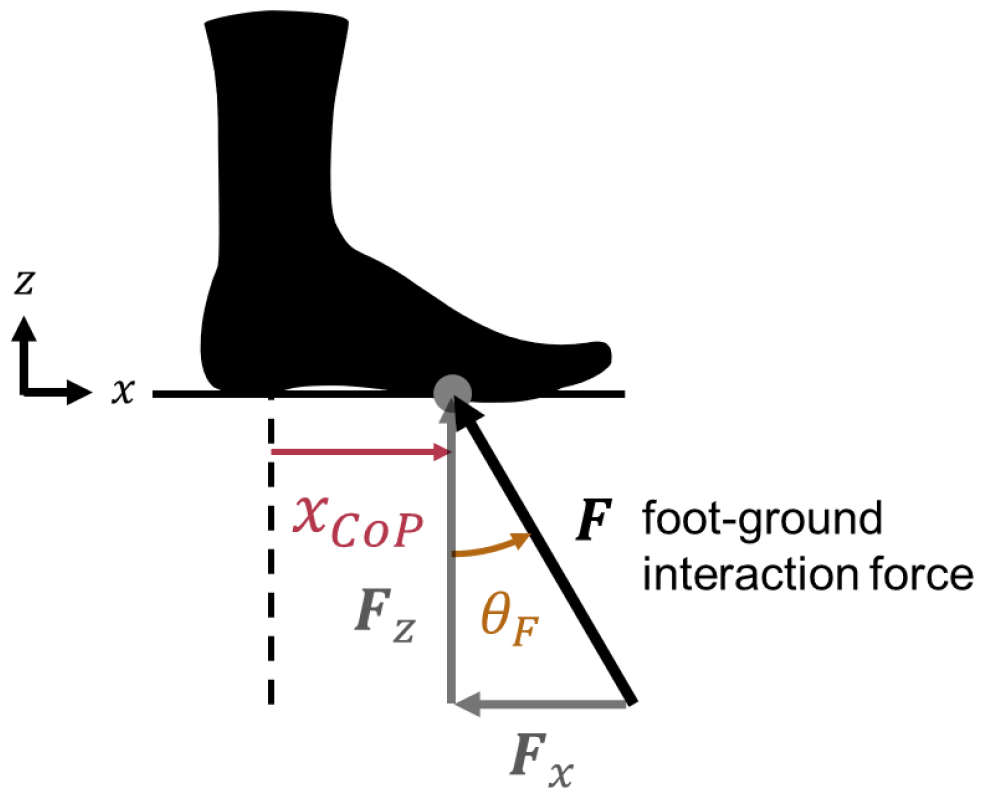
Illustration of the center-of-pressure position (*x*_*CoP*_) and force angle (θ_*F*_) signals with respect to the foot-ground interaction force vector (***F***). Vertical (***F***_*z*_) and horizontal (***F***_x_) force components are shown as well. The center-of-pressure position was defined with respect to the origin of the force-plate coordinate system.

First, the θ_*F*_ and *x*_*CoP*_ time-series signals were detrended. Then, the 2-by-2 cross-spectral density matrix between these two signals was computed, and the real component, the co-spectral density matrix, was extracted. The height of the intersection point at each frequency band was the slope of the principal eigenvector of the co-spectral density matrix of *x*_*CoP*_ versus θ_*F*_. The intersection-point height could be related to Δ*x*_*CoP*_, small changes in the frequency-filtered center-of-pressure position, and Δθ_*F*_, small changes in the frequency-filtered foot-force orientation, by

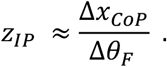

The intersection-point height for each trial of each participant was obtained in the sagittal plane, using the anteroposterior component of the center of pressure (*x*_*CoP*_) and the foot-force orientation (θ_*F*_). The spectral method was implemented with a frequency resolution of 0.098 Hz. The four lowest-frequency intersection-point-height estimates were discarded due to artifacts from data pre-processing (23), and estimates up to 8 Hz were considered in the final analysis. This choice gave a final frequency range of 0.49 – 7.91 Hz. The intersection-point height at each frequency was normalized by the center-of-mass height of each participant and averaged across the three trials to find the per-participant intersection-point height, and then further averaged across the participants within each age group to find the average and standard deviation for each group.

### Center-of-Pressure Speed

Having tested 12 different center-of-pressure-based measures of quiet standing, Prieto et al. identified mean center-of-pressure speed as the measure that showed the greatest correlation with age (14). Thus, mean center-of-pressure speed was used as a comparison to evaluate the effectiveness of the intersection-point measure in differentiating between age groups.

As the intersection-point measure was computed in the sagittal plane, the mean center-of-pressure speed was also computed for the anteroposterior (AP) axis and denoted as 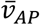.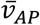 was computed by summing the length of the center-of-pressure path along the anteroposterior axis (sum of displacements between adjacent center-of-pressure sample points) and dividing that by the total time, *T*:

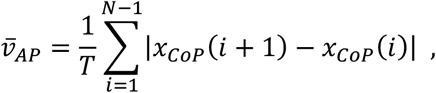

where *N* was the number of time samples. The mean center-of-pressure speed was computed for each trial of each participant, and then averaged over the three trials per participant to obtain a single speed measure per participant. The mean for each age group was also computed by averaging across participants.

## Modeling

### Biomechanical Model

Though still commonly used in balance studies (24–26), a single-inverted-pendulum model could not replicate the intersection point below the center-of-mass height that was observed in humans (11). As shown in Fig. 2, the next simplest model, a double-inverted-pendulum model, was employed.

**Figure 2.**
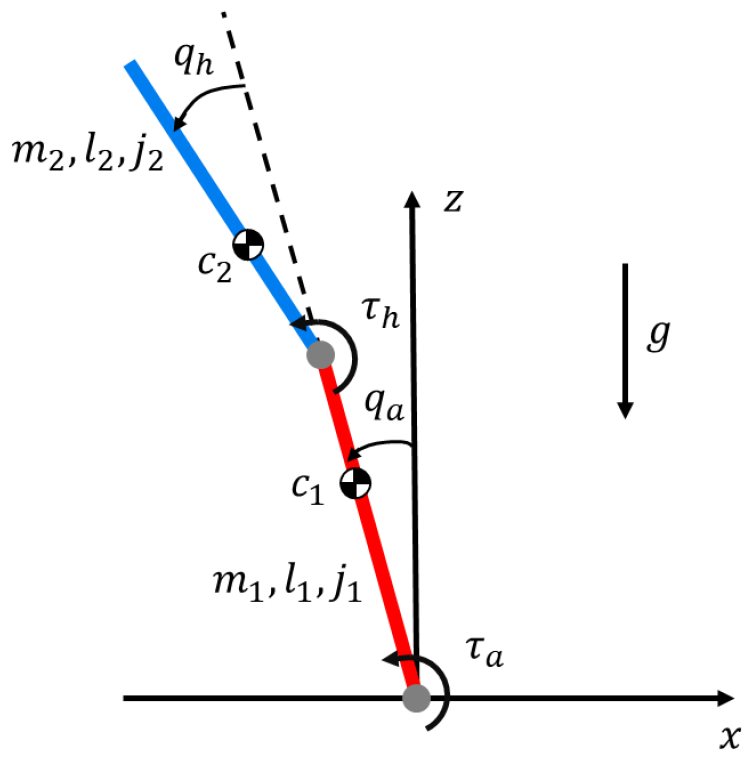
Double-inverted-pendulum model in the sagittal plane. The bottom red segment represents the lower body and the blue link above it reflects the upper body. The labels indicate angle (*q*_*a*_, ankle; *q*_*h*_, hip) and torque (τ_*a*_, ankle; τ_*h*_, hip) conventions and parameter values for mass (*m*_*i*_), length (*l*_*i*_), center-of-mass position (*c*_*i*_), and moment of inertia about the center of mass (*j*_*i*_). τ_*a*_ is the torque acting on the lower body from the ground, while τ_*h*_ is the torque acting on the upper body from the lower body. The direction of gravity (*g*) is also illustrated.

Based on the anthropometric distribution of young male participants (27) and the average mass and height of the older and younger participants, the lumped model parameters were determined for each participant group (Table 1). Preliminary analysis showed no differences in outcomes when using the anthropometric distribution of female participants instead. Because the foot was stationary, any mass and height below the ankle were neglected. The foot being stationary also implied that the sum of the torques on the foot were equal to zero; from this torque balance and from the height of the foot being zero, the center of pressure was computed as:

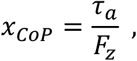

where *F*_*z*_ was equal to the force of gravity plus the vertical acceleration of the center of mass times the body mass, as defined by Shiozawa et al. (11). The center-of-mass positions for each link were measured with respect to the ankle joint for the lower body (link 1) and the hip joint for the upper body (link 2).

**Table 1.** Lumped model parameters.

| Symbol | Parameter (units) | Older Participants Value |  | Younger Participants Value |  |
| --- | --- | --- | --- | --- | --- |
|  |  | Link 1<br>Lower Body | Link 2<br>Upper Body | Link 1<br>Lower Body | Link 2<br>Upper Body |
| $m$ | Mass (kg) | 23.33 | 38.04 | 23.15 | 37.74 |
| $l$ | Length (m) | 0.778 | 0.763 | 0.832 | 0.817 |
| $l_c$ | Center of mass (m) | 0.528 | 0.342 | 0.565 | 0.366 |
| $j$ | Moment of inertia (kgm<sup>2</sup>) | 1.000 | 1.490 | 1.136 | 1.693 |
| $g$ | Gravitational acceleration (m/s<sup>2</sup>) | 9.81 | | | |

The moments of inertia were calculated about the center of mass of each link. The equations of motion and the foot-force vector orientation, which were also required to compute the intersection-point height above the floor, were calculated in a similar manner as in a previous work (11). The state and input variables were 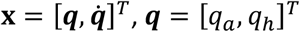, and **τ** = [τ_*a*_, τ_*h*_]^*T*^ as defined in Fig. 2.

Noise processes were added to the ankle and hip joint torques to model the stochastic nature of the biological controller and were assumed to be white, mutually uncorrelated, and normally distributed (with zero mean). *σ*_*ankle*_ and *σ*_*hip*_ corresponded to standard deviations of the noise on the ankle and hip torques. The relative strength of the two noise sources was defined as 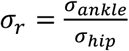.

### Controller

The linear quadratic regulator (LQR), which guarantees a stable closed-loop system for all design parameter values, was used. The nonlinear equations of motion for the double-inverted pendulum were first linearized about the upright posture **x**_0_ = [0, 0, 0, 0]^*T*^, **τ**_0_ = [0, 0]^*T*^. The LQR computes an optimal control gain matrix, ***K***_*LQR*_ ∈ ℝ_2×4_, based on the quadratic cost function

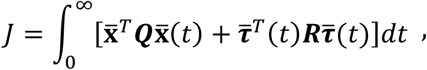

where 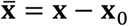 and 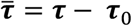, and ***Q*** and ***R*** weight the state and control-input deviations, respectively, from zero. This cost function is minimized to determine the control torques

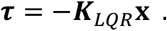

The control-input weighting matrix was parameterized as

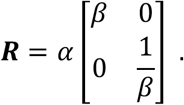

with design parameters *α* and *β*. When *α* was set to be large, the closed-loop system exhibited a predetermined behavior by placing its poles at the mirrored positions of the unstable open-loop poles; this behavior was unaffected by the specific choice of the state weighting matrix ***Q***, as long as ***R*** was sufficiently larger than ***Q***. The effect of *α* on the intersection point was explicitly analyzed previously (11), and large *α* was shown to reduce the control effort to a minimum value required for stability and to best fit healthy, young human data. Proceeding with the assumption that unimpaired balance is best described by a controller that minimizes effort, *α* was set to be large (10^6^) for this study. This choice also reduced the number of control parameters to search through to just one (*β*). When *β* > 1, the ankle torque was penalized more heavily than the hip, and vice versa when *β* < 1. Although the poles did not vary significantly for large *α*, the closed-loop gain matrix varied considerably with *β*.

### Best-Fit Control Parameter Search

To quantify the variation of best-fit parameters for the older and younger age groups respectively, parameters were varied systematically. An analytic approach (23) was used to compute the intersection-point height normalized by the center-of-mass height and facilitated faster iteration through the *β* and σ_*r*_ parameters. Because human balance evokes only small vertical motions, a constant center-of-mass height based on the participant group’s average height was assumed.

#### Best-Fit Parameters for Average z_IP_

To search for the parameter set that fit the average intersection-point heights for each age group, the following procedure was employed. First, the parameter that weights the relative cost of the control input, *α*, was set to 10^6^. Then, a global parameter search of the parameter that adjusts the relative cost on the hip and ankle torques, *β*, and the joint actuation noise ratio, σ_*r*_, was conducted. For *β*, values between 0.01 and 3 were explored; for σ_*r*_, values between 0.01 and 20 were explored.

To further determine age-related differences, a finer search was conducted. For *β*, values between 0.15 and 0.45 were explored; for σ_*r*_, values between 0.01 and 1 were explored. These narrower ranges were selected based on a preliminary analysis evaluating regions with the highest concentration of best-fit parameters on individual intersection-point data.

#### Best-Fit Parameters for Per-Participant z_IP_

The finer parameter search was also conducted to find the best-fit parameters for each participant’s frequency-dependent intersection-point-height curve. Similar to above, *α* was set to be 10^6^, while values between 0.15 and 0.45 were explored for *β*, and values between 0.01 and 1 were explored for σ_*r*_.

#### Comparison of Experimental and Simulated Results

To determine the best-fit intersection-point-height curve across different model parameters, the rootmean-squared error (RMSE) between the computed and experimental data was calculated over all frequency bands by

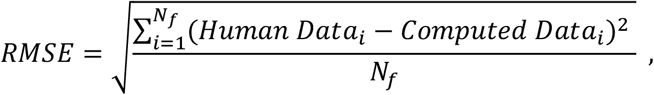

where *Human Data*_*i*_ was the average observed intersection-point height (either across participants for each age group, or across all trials for each participant). *Computed Data*_*i*_ was the analytically computed intersection-point height (23) in a given frequency band. *N*_*f*_ was the number of frequency bands in which the intersection-point height was computed: 77 for this study. The best-fit parameter set was the one with the lowest *RMSE* value across all tested *β* and σ_*r*_ values.

### Optimal Gains

For each best-fit control parameter set, the optimal gain matrix ***K***_*LQR*_ = [***K, B***] with stiffness, ***K*** ∈ ℝ_2×2_, and damping, ***B*** ∈ ℝ_2×2_, components were recorded. Any square matrix can be partitioned into a symmetric component and an antisymmetric component. For example, the stiffness matrix can be rewritten as:

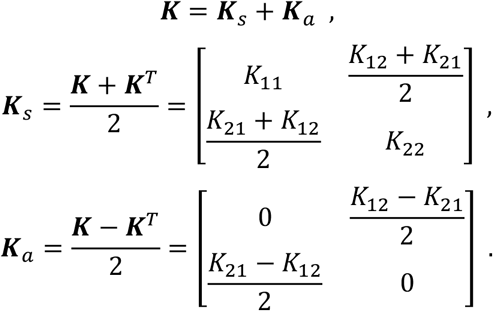

where ***K***_*s*_ and ***K***_*a*_ are the symmetric and antisymmetric components of ***K***, and *k*_*ij*_ represents the element in the *i*th row, *j*th column of ***K***. The eigenvalues and eigenvectors of the symmetric and antisymmetric components were computed to illustrate the matrices graphically. The symmetric component was represented as an ellipse: the major axis length was determined by the principal eigenvalue, while its orientation was determined by the corresponding eigenvector; the second largest eigenvalue and eigenvector similarly determined the length and direction of the minor axis of the ellipse. The antisymmetric component was illustrated as a circle: its radius was equivalent to the absolute value of the eigenvalue.

Previous work has shown that while the symmetric component of stiffness has conservative properties, the antisymmetric component constitutes forces that cannot be derived from a potential function (28, 29). Notably, a non-zero antisymmetric component of stiffness can only be attributed to neural feedback and not to the biomechanical properties of muscles. While the appearance of an anti-symmetric component of stiffness can unequivocally be assigned to neural feedback, neither the symmetric nor the anti-symmetric components of damping may unequivocally be assigned to neural feedback. For that reason, further analysis focused on stiffness.

Comparing the relative magnitudes of the symmetric and antisymmetric components of stiffness enabled the evaluation of the extent to which the closed-loop gain matrix can be attributed to neural feedback (30). To this end, their ratio was computed as follows:

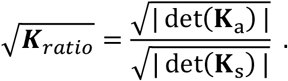

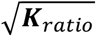 can range from 0 to ∞, where a value of 0 corresponds to a purely spring-like neuromuscular system, while 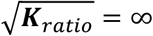 can result from a purely rotational field due to neural feedback. This ratio was computed for the closed-loop stiffness matrix based on both the average and per-participant bestfit controller parameters. Note that although the symmetric and antisymmetric components of the stiffness matrix may change with a different choice of coordinates, their ratio is unaffected. Since 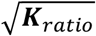 is coordinate independent, it served as an appropriate tool to compare the role of feedback in older and younger participants.

### Simulation Protocol

Once the best-fit parameters were determined for the average intersection-point height of each participant group, 40 simulation trials were conducted to enable statistical analysis of the simulated dependence of the intersection-point height on frequency. The simulation was conducted using semi-implicit Euler integration. The initial condition was set to **x**_0_. Each simulation was run for 60 s at 1000 Hz, and the output data were down-sampled to 100 Hz to replicate the conditions of the human experiments. A constant center-of-mass height based on the participant group’s average height was assumed to compute the intersection-point height normalized by the center-of-mass height. All simulations were conducted in MATLAB 2022b (MathWorks, Natick MA). Note that the analytic approach yielded essentially identical results to the numerically-computed intersection-point height (23).

The control input at each time step for each simulation trial was stored to compute the root-mean-squared (RMS) value of the torque for the ankle and hip joints separately. Then, to compare the torque contributions from each joint, the ratio of the joints’ RMS torques was computed as

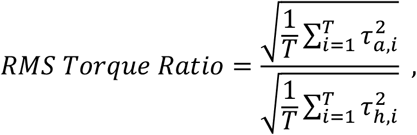

where τ_*a*_ and τ_*h*_ are ankle and hip torques, and *T* is the number of time steps. Finally, across the 40 trials for the best-fit *β* and σ_*r*_ parameters, the mean and standard deviation of the RMS torque ratio were reported.

### Statistical Analysis

To determine whether certain measures could distinguish between older and younger participants, statistical analysis was conducted. Because of the large sample size, the sample means were assumed to be normally distributed (31). All statistical analyses were carried out in MATLAB 2022b. The significance level was set to *p* = 0.05 for all statistical tests.

#### Intersection Point

To test whether the mean intersection-point-height curves of the two age groups were significantly different, the difference between the two curves at each of the 77 frequency points were computed, and the distribution of the differences was analyzed. A one-sample t-test was applied to determine whether the mean difference between the curves was significantly different from zero.

#### Center-of-Pressure Speed

To test whether the mean anteroposterior center-of-pressure speeds of the two age groups were significantly different, Welch’s t-test was conducted.

#### Best-Fit Controller Parameters

After obtaining the best-fit controller parameters for each participant’s frequency-dependent intersection-point curve, Welch’s t-test was conducted for *β* and σ_*r*_ parameters separately to determine the effect of age.

#### Optimal Stiffness

After obtaining 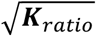 for each participant, values that were three standard deviations away from the mean (two samples from the younger participants’ data, 2% of the entire dataset) were excluded. Then, Welch’s ttest was conducted to determine the effect of age.

## RESULTS

### Human Experiment Results

#### Intersection-Point Height

The frequency-dependent intersection-point-height pattern for older and younger participants exhibited similar trends where the intersection point was above the center of mass at low frequencies and below the center of mass at high frequencies (Fig. 3). However, specific details differed between the two groups. For example, the mean of the intersection-point height was approximately equal to the center-of-mass height at 2.1 Hz and 1.8 Hz for older and younger participants, respectively. At 4.4 Hz, the mean of the intersection-point height was 0.61 for older participants and 0.33 for younger participants. The distribution of the difference between the mean intersection-point height of older and younger participants was greater than zero and significantly different from a zero-mean normal distribution (*p* = 3.5 × 10^−34^ < 0.05), indicating that the mean intersection-point height curve of older participants was significantly higher than that of the younger participants.

**Figure 3.**
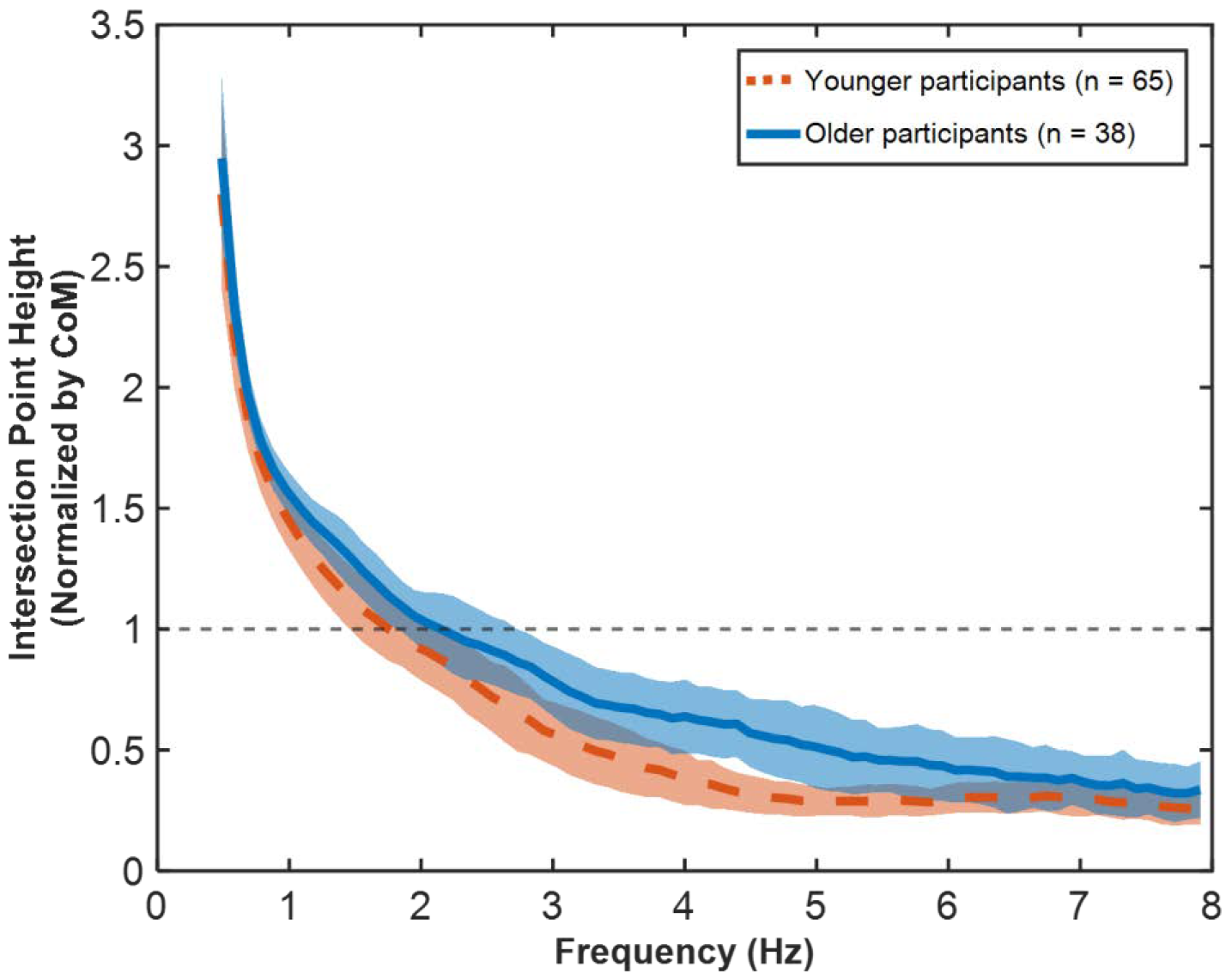
Older (n = 38) and younger (n = 65) participant data of mean intersection-point heights normalized by the center-of-mass height as a function of frequency. The shaded areas indicate one standard deviation of the data. The center-of-mass height was computed for each participant. The normalized height of the center of mass is marked by a dashed line.

#### Center-of-Pressure Speed

The mean anteroposterior center-of-pressure speed averaged over all the older participants was 9.0 ± 3.9 mm/s (mean ± standard deviation), while that of the younger participants was 6.5 ± 2.0 mm/s. Welch’s t-test showed a significant effect of age (*p* = 1.0 × 10^−9^ < 0.05).

### Best-Fit Controller Parameters

#### Global Parameter Search

When a global parameter search was conducted assuming minimal control effort (*α* = 10^6^), the best-fit parameter set that achieved a good fit at all frequencies was *β* = 0.2 and σ_*r*_ = 0.6 for older participants and *β* = 0.3 and σ_*r*_ = 0.7 for younger participants. These best-fit parameters yielded fits with a RMSE of 0.1536 for the older participant group and 0.1384 for the younger participant group. As shown in Fig. 4a and 4b, there is a narrow range of *β* and σ_*r*_ parameters that produced a good fit of the human data for both participant groups. In particular, smaller RMSE values were observed for *β* values between 0.15 and 0.45 and σ_*r*_ values between 0.01 and 1. To explore those parameter ranges further, a finer search was conducted in those regions.

**Figure 4.**
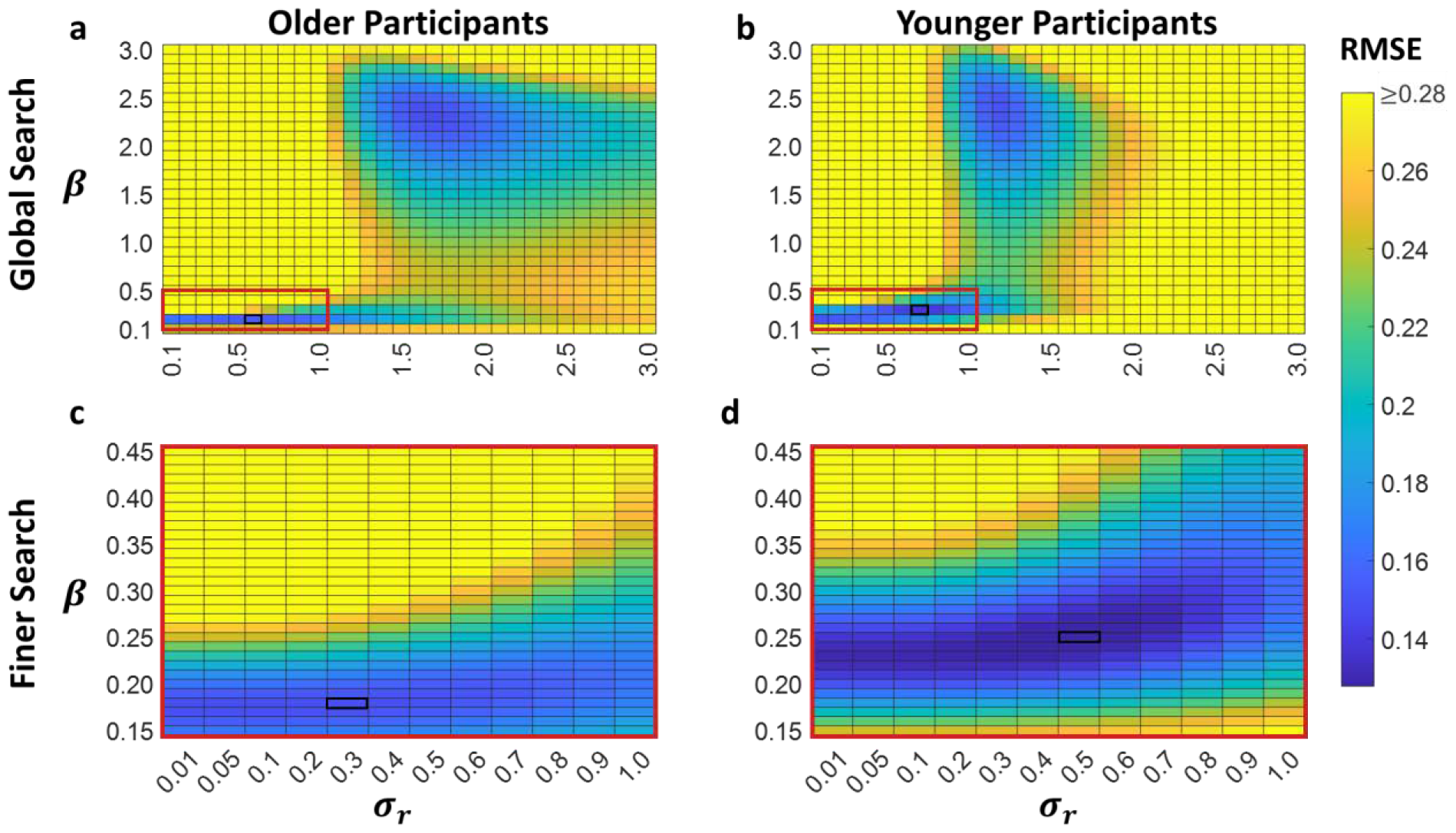
RMSE across a range of varied parameters for the two participant groups: a) global parameter search on older participants’ data, b) global parameter search on younger participants’ data, c) finer parameter search on older participants’ data, d) finer parameter search on younger participants’ data. The red rectangles in the global search plots indicate the range of parameters that the finer search covered. The black rectangles in all of the plots indicate *β* and σ_*r*_ value sets that yielded the smallest RMSE for each search. The color bar indicates the RMSE values for all subplots. All RMSE values greater than or equal to 0.28 were represented by the same shade (yellow). Some tested *β* and σ_*r*_ values were not reported in this graphic to maintain the continuity of the two axes.

#### Finer Parameter Search

When a finer parameter search was conducted, the best-fit parameter set was *β* = 0.18 and σ_*r*_ = 0.3 for older participants and *β* = 0.25 and σ_*r*_ = 0.5 for younger participants. These parameter sets resulted in a RMSE of 0.1488 for the older participants and 0.1245 for the younger participants, which was overall smaller than the results of the global parameter search (Fig. 4c and 4d). With the best-fit parameter sets from the finer search, the frequency-dependent intersection-point-height curves were reproduced well for both older and younger participants, as shown in Fig. 5. These best-fit parameters resulted in RMS torque ratios of 3.91 ± 0.02 (mean ± standard deviation) and 2.07 ± 0.02 for older and younger participants, respectively, indicating that older participants typically used ankle joint torque to a greater extent than younger participants.

**Figure 5.**
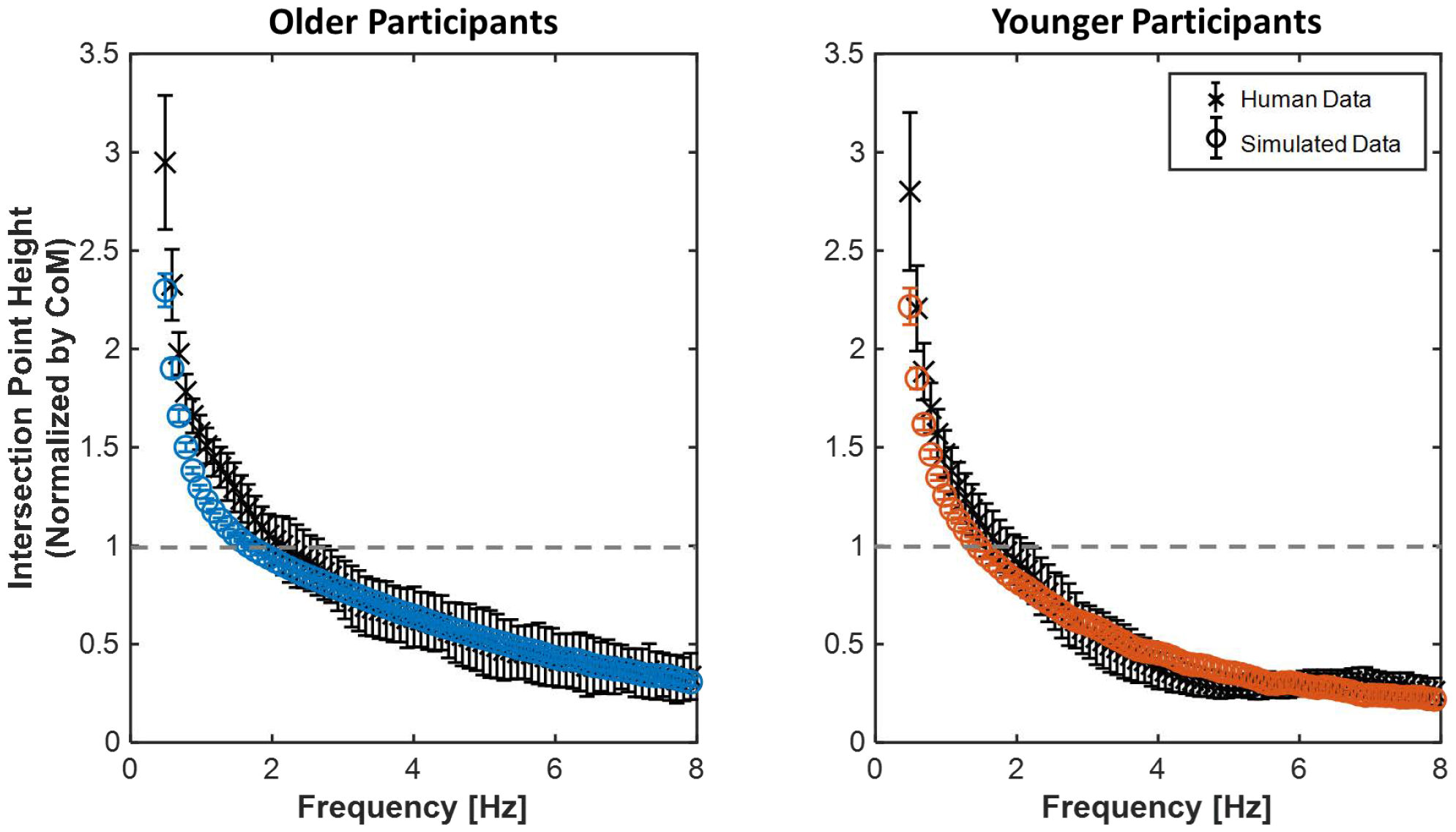
Mean intersection-point heights normalized by the center-of-mass height of older participants (left) and younger participants (right). Human data are shown with x, and the mean of the best-fit simulation results are shown with o. The error bars denote one standard deviation across the participants for the human data and across 40 simulation runs for the simulated data. The height of the center of mass is marked by a dashed line. For the human data, the center-of-mass height was computed for each participant; for the simulated data, the center-of-mass height was calculated based on the average height of the participants in each age group and kept constant.

#### Per-Participant Best-Fit Controller Parameters

A fine search of best-fit parameters for each tested participant resulted in means of *β* = 0.19 ± 0.01 and σ_*r*_ = 0.34 ± 0.12 for older participants and *β* = 0.26 ± 0.01 and σ_*r*_ = 0.42 ± 0.06 for younger participants (all with 95% confidence), as shown in Fig. 6. The best-fit *β* parameter in older participants was statistically different than the best-fit *β* parameter for younger participants (*p* = 1.6 × 10^−15^ < 0.05). This result indicates that older participants penalized the use of the hip more than the younger participants. On the other hand, σ_*r*_ did not show a significant difference between the two participant groups (*p* = 0.23 > 0.05).

**Figure 6.**
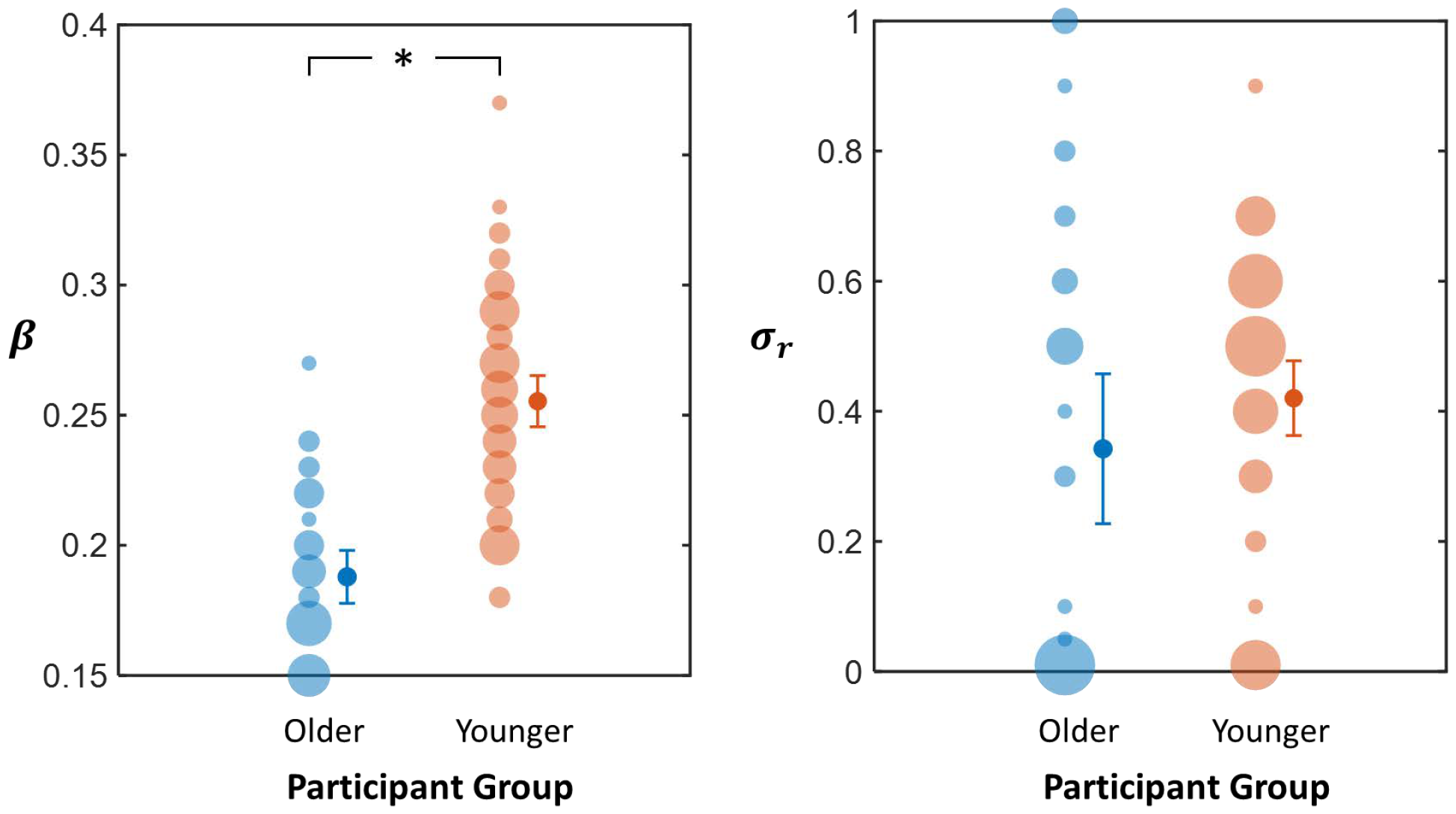
Distribution of *β* (left) and σ_*r*_ (right) parameter values that best fit the older and younger participants’ individual data. The size of the transparent circles represent the number of datapoints; larger circles had more datapoints corresponding to a particular value. The mean and 95% confidence interval are shown to the right of each group’s distribution, and the values were *β* = 0.19 ± 0.01 and σ_*r*_ = 0.34 ± 0.12 for older participants and *β* = 0.26 ± 0.01 and σ_*r*_ = 0.42 ± 0.06 for younger participants. While the *β* parameter showed statistical difference between the older and younger participants (*p* = 1.6 × 10^−15^ < 0.05), σ_*r*_ did not (*p* = 0.23 > 0.05).

### Best-Fit Stiffness

The best-fit controller parameters for the average intersection-point curves of each participant group yielded the following stiffness gains:

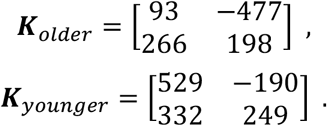

A graphical representation of the symmetric and antisymmetric components of the stiffness matrices for each participant group is shown in Fig. 7.

**Figure 7.**
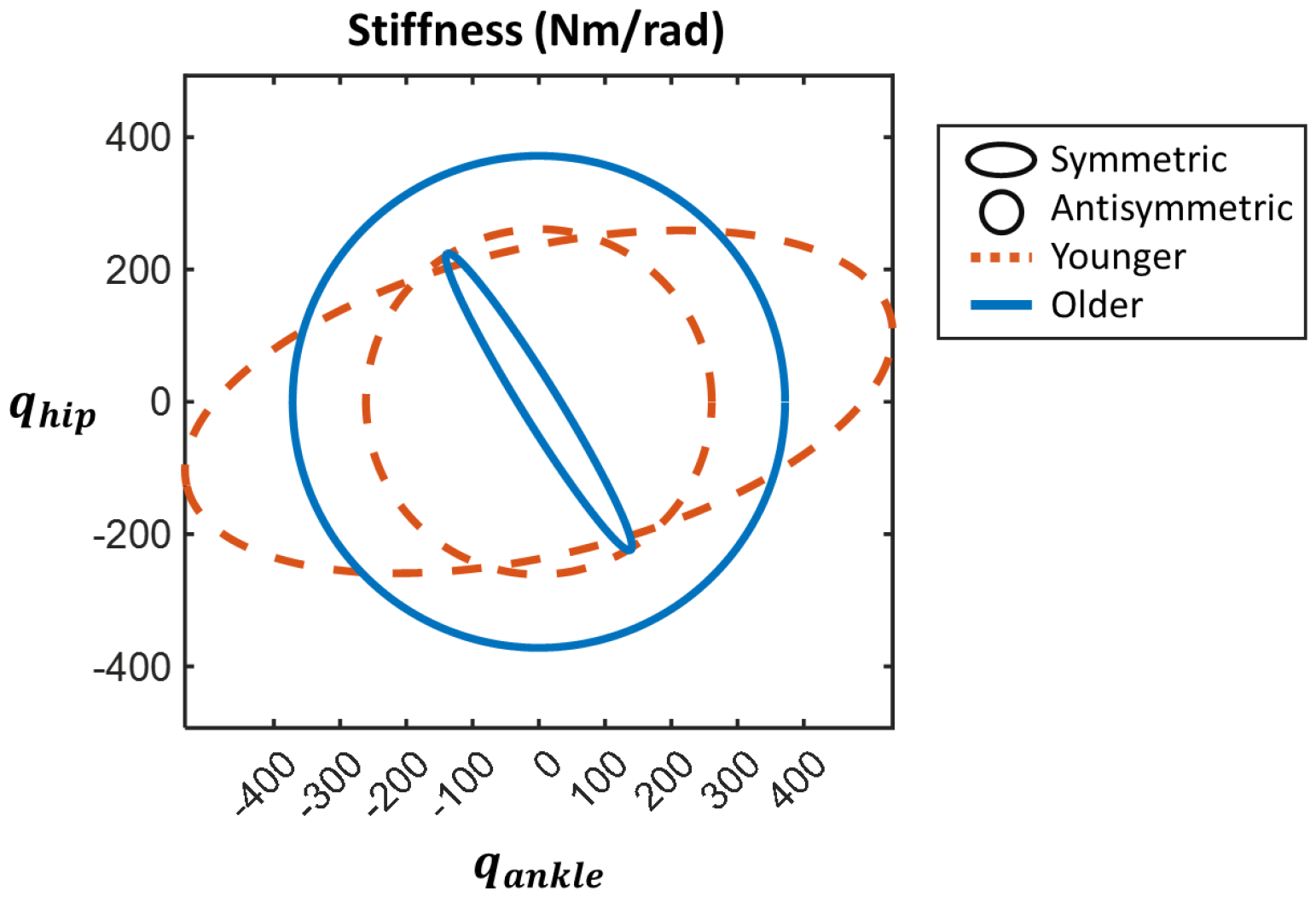
Graphical representation of the symmetric and antisymmetric components of the stiffness matrices based on the best-fit control parameter for each participant group. The blue solid lines correspond to older participants, while the orange dotted lines correspond to younger participants. The symmetric component was represented as an ellipse: the major axis’ length was determined by the principal eigenvalue, while its orientation was determined by the corresponding eigenvector; the second largest eigenvalue and eigenvector similarly determined the length and direction of the minor axis of the ellipse. The antisymmetric component was illustrated as a circle; its radius was equivalent to the absolute value of the matrix’s eigenvalue.

When the relative magnitude of the symmetric and antisymmetric components were compared, the 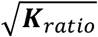 was 4.33 for older participants and 0.73 for younger participants. The relative magnitude of the symmetric and antisymmetric components was also computed for each participant’s best-fit stiffness gains. The mean 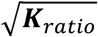 was 1.34 ± 0.29 for older participants and 0.79 ± 0.08 for younger participants with 95% confidence, as summarized in Fig. 8. Older participants’ 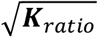 was statistically greater than those of younger participants (*p* = 6.7 × 10^−4^ < 0.05).

**Figure 8.**
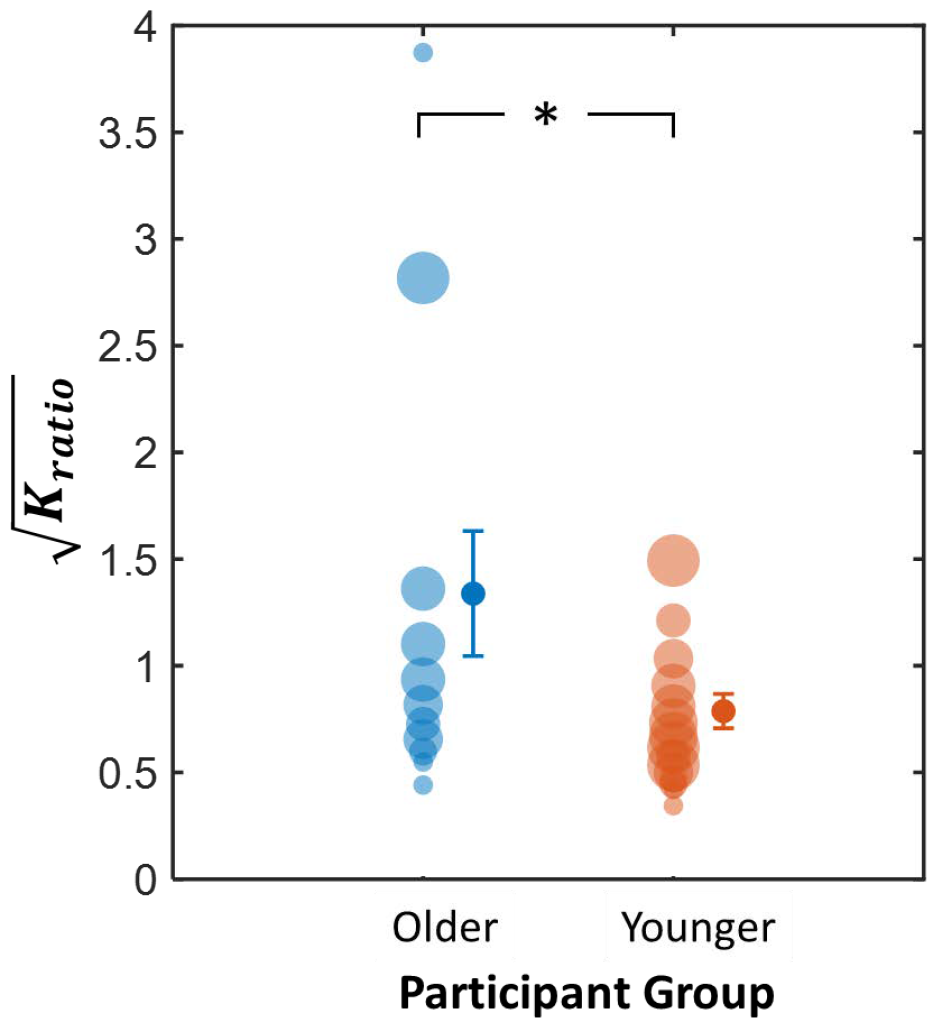
Distribution of the 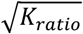 based on the best-fit stiffness matrices for older and younger participants’ individual data. The size of the transparent circles represent the number of datapoints; larger circles had more datapoints corresponding to a particular value. The mean and 95% confidence interval are shown to the right of each group’s distribution, and the values were 1.34 ± 0.29 for older participants and 0.79 ± 0.08 for younger participants. There was a statistical difference between the older and younger participants (*p* = 6.7 × 10^−4^ < 0.05).

## DISCUSSION

This study compared balance control strategies between healthy younger and older adults by analyzing a previously published dataset in which 65 younger participants and 38 older participants were tasked to stand quietly (22). Two measures, the sagittal-plane intersection point of foot-ground forces and mean anteroposterior center-of-pressure speed, were obtained from the human data. An intersection-point measure was able to differentiate between age groups at least as well as the traditionally-employed center-of-pressure measure, but in addition, the intersection-point analysis enabled quantification of controller parameters that described the human data using a simple nonlinear human balance model with a linear optimal controller. This method revealed age-specific control strategies that suggested greater reliance on neural feedback in older participants. These findings demonstrate the potential of intersection-point analysis as a quantitative clinical assessment tool to measure the state of the balance controller.

### Intersection-Point Height vs. Center-of-Pressure Speed

The competency of both the intersection-point height and the mean center-of-pressure speed to distinguish between the age groups was first evaluated. Both measures demonstrated statistically significant differences between older and younger participants. These results indicated that, like the mean center-of-pressure speed, a traditionally employed measure, the intersection point could differentiate age groups.

Detailed findings revealed that the intersection point was on average higher in older participants compared to younger participants, particularly in the middle to higher range of frequencies where the intersection point was near the knee (Fig. 3). An intersection point at the ankle would correspond to control solely by hip torque if balance is modeled by a double-inverted pendulum (11). Thus, the observation that older participants presented with heights that are further away from the ankle in the mid to high frequency ranges indicates decreased actuation of the hip relative to the ankle. Also, the older participants had faster center-of-pressure speeds compared to younger participants, aligning with prior research and suggesting increased activity of the postural control system in older adults (14).

Although both the intersection-point height and the center-of-pressure speed could afford insight into the differences between the two age groups, the incorporation of the foot-ground force orientation in the intersection-point measure adds valuable information about the dynamics of postural control. Moreover, by employing a two-segment model and applying it to the intersection-point measure, we successfully quantified distinct control strategies that best described the human data for each age group, as summarized in the next section.

### Best-fit Parameters

Comparing the intersection-point height of human data with simulation outcomes facilitated the identification of age-specific control strategies. Both global and finer searches for the best-fit parameter set indicated a higher hip penalty or a more pronounced use of the ankle in older participants compared to younger participants. The optimal parameters were fairly narrowly defined, especially for younger participants, demonstrating that the results were less likely to be vulnerable to small changes in assumptions. Per-participant results showed a significant difference across the two participant groups for the *β* parameter, suggesting a variation in the relative use of the ankle and hip joints across age groups. Previous research showed that increased ankle stiffness counteracted destabilizing forces in systems with heightened neural delay (32). As response to perturbation can be delayed by 10 to 30 ms in older participants compared to younger participants (33, 34), older participants may have compensated by increasing their use of the ankle.

The analysis also revealed that the noise in the hip was larger than the noise in the ankle for both age groups. This result is consistent with a prior study on young, unimpaired participants tasked to balance with feet shoulder-width apart (11). Furthermore, in a preliminary study of a global search of the best-fit parameters for each participant, a trend emerged: when the ankle was noisier than the hip, ankle torque was penalized more, and vice versa. This finding suggests a general tendency in humans balancing quietly with feet shoulder-width apart: in the sagittal plane, larger noise in a joint led to a greater penalty on that joint.

### Stiffness

We additionally analyzed the stiffness matrix of the closed-loop system, using the best-fit controller values derived from both the average and individual data. The focus was on discerning differences between the symmetric and antisymmetric components of the stiffness matrix, as they quantify different roles of control (28). The antisymmetric component of stiffness represents forces with curl, not derivable from a potential function, while the symmetric component has conservative properties.

Physiologically, the symmetric component quantifies both intrinsic muscle stiffness and neural feedback gains, whereas the antisymmetric component is non-zero only if inter-segmental feedback action exists in the limbs.

The study’s results revealed that both older and younger participants exhibited a non-zero rotational or curl component of stiffness, consistent with the significant role of feedback in maintaining balance. This observation aligns with previous studies that concluded that intrinsic muscle stiffness in the ankle alone is insufficient to maintain stability during upright stance (35, 36).

Moreover, upon comparing the relative magnitudes of the antisymmetric and symmetric components of stiffness for each participant, the mean of the distributions revealed that older participants had statistically larger antisymmetric components compared to their symmetric components, while younger participants exhibited the opposite behavior. This finding suggests diminished intrinsic muscle stiffness in older adults, which is potentially indicative of muscle strength loss or sarcopenia. Agreeing with this finding, previous work found reduced structural stiffness in the triceps surae, tibialis anterior, quadriceps, and hamstrings in older adults (37–39). Some of these studies further found that decreased muscle stiffness was related to lower muscle mass and increased rate of sarcopenia (38, 39).

Remarkably, the larger antisymmetric component observed in older adults suggests a compensatory use of neural feedback to offset this decline in strength and/or stiffness.

### Beyond Age

The aforementioned results highlighted a statistically significant increase of the intersection-point height and penalty on the hip joint among adults over the age of 60. However, a closer examination of all individual participant data revealed that there was a sizable variation in the control parameter that determined the relative cost of control effort in the ankle compared to the hip (*β*) even within age groups. This finding emphasizes that age is not the only factor influencing balance control. Similarly, a previous study did not find variations in mean center-of-pressure speed with age in hospitalized older participants, as their balance impairments were likely dominated by underlying pathologies (40).

Each individual may carry unique, underlying pathologies that contribute to balance deficits irrespective of their age (41). Currently, balance impairment diagnosis relies on clinical assessments, such as the MiniBESTest or the Berg Balance Test. Because the intersection-point analysis showed variation of postural control strategies beyond age-related differences, it may provide an additional factor of analysis to better inform clinicians of the degree of a patient’s postural dyscontrol.

### Quantitative Clinical Assessments

As considering individualized factors is important in the evaluation of postural imbalance, there is a pressing need for quantitative clinical assessments that are capable of precisely articulating the extent of an individual’s balance impairment. Current clinical assessments often rely on qualitative methods, and results can vary based on the therapist’s subjective observations. While informal observation-based assessments remain prevalent (42), many physiotherapists acknowledge the importance of incorporating objective quantitative measures (42, 43). However, clinicians are dissatisfied with widely-used quantitative balance assessments, such as the Single-Leg Stance, Berg Balance, and Timed “Up and Go” tests, citing concerns about their time-intensive nature, limited applicability, and lack of comprehensiveness (42, 43).

In this context, the proposed intersection-point analysis offers a valuable alternative. This method can quantify a patient’s deviation from typical balance control in less than one minute. With its ability to provide quantitative insights on control strategy, the intersection-point analysis also has the potential to enhance clinicians’ understanding of the underlying causes of impairment. This shift towards quantitative, detailed assessments aligns with the evolving needs of clinicians seeking a more nuanced understanding of balance impairments for improved treatment strategies.

### Limitations

Using a simplified model inevitably introduces limitations. The chosen model incorporated two degrees of freedom, the minimum required to reproduce the experimental data (11). This model excluded the foot’s height and mass for the computation of the center-of-pressure position and center-of-mass height. Preliminary analysis showed that this assumption had little effect on the results. For the model’s two degrees of freedom, we selected relative angles as the coordinate frame. Therefore, the reported stiffness matrices are represented in those coordinates. The age-related differences in the role of the feedback was evaluated based on a coordinate-independent measure given by the ratio of the matrix determinants; however, the specific form of the stiffness matrices and their symmetric and antisymmetric components must be interpreted with care. The model also used the anthropometric body-mass distribution based on young adults (27), which may not represent the older participants considered in the data set (22). Future work should investigate the sensitivity of the intersection-point-height measure to the biomechanical model parameters.

Additionally, while the biomechanical model employed was nonlinear, the controller design relied on linearization. Consequently, the best-fit parameters reported in this study are valid primarily for small motions near the upright position. This assumption was appropriate, as experimental data consisted of participants standing quietly on a stationary surface, where larger body motions were not expected.

Moreover, the assumption that humans minimize their control effort reduced the search space to two parameters. Although we focused on designing the simplest descriptive model and controller, other formulations may also be able to account for the data.

The simulations conducted in this work assumed highly simplified neuro-mechanics. Notably, factors such as neural transmission delay and sensory noise were not explicitly included in the model. The ankle and hip torques summarized the action of multiple muscles. These simplifications allowed for the application of the proposed method to participants across a broad age range. Future work should implement this model to understand human data of persons with neural impairments to investigate control strategies underlying the decline of balance ability.

## CONCLUSION

The intersection-point measure can be more informative than traditionally applied balance measures, as it incorporates the orientation of the foot-ground force in addition to its point of application. In this study, simulating this intersection point during quiet balance using a simple, interpretable model and comparing the results to human data enabled the identification of distinct control strategies in neurologically unimpaired older and younger participants. A closer examination of the best-fit optimal control gains suggested increased reliance on neural feedback in older participants that possibly compensated for muscle degradation and increased neural transmission delay. This work further demonstrated the intersection-point analysis’ ability to be applied to individual participant data, holding promise for the development of future quantitative diagnostic tools. To broaden the applicability of the findings and tools of this work, further investigation is needed to analyze individuals with neural impairments, such as stroke survivors.

## DATA AVAILABILITY

Source data for this study were derived from Santos and Duarte (2016), available at https://dx.doi.org/10.6084/m9.figshare.3394432.v2.

ACKNOWLEDGMENTS

None.

## DISCLAIMERS

None.

## AUTHOR CONTRIBUTIONS

NH and KG contributed to conceiving and designing the research, RSD processed human experimental data, KS performed simulation experiments and processed simulation data. All authors contributed to interpreting the results of experiments. KS and RSD drafted the manuscript, KS prepared the figures, and all authors edited, revised, and approved the final version of the manuscript.

